# Laboratory sprayer for testing of microbial biocontrol agents: design and calibration

**DOI:** 10.1101/2020.04.22.054551

**Authors:** Z. Erdos, P. Halswell, A. Matthews, B. Raymond

**Affiliations:** University of Exeter, Penryn TR10 9FE, United Kingdom

**Keywords:** biological control, spray tower, entomopathogen fungi, *Beauveria*, *Akanthomyces*, Myzus persicae

## Abstract

The lack of commercially available low-cost laboratory spraying equipment for testing microbial control agents can hinder advancement in the field of biocontrol. This study presents an inexpensive, portable sprayer that is calibrated utilizing laboratory consumables. The computer aided design files are made available so that it is freely modifiable and can be used for machine routing or 3D printing. Bioassay data was obtained by spraying *Myzus persicae* with spores of entomopathogenic fungi. Observed variation in droplet deposition within tested pressure and volume settings, and spore deposition within sprayed concentrations were low. Bioassay results show reproducible mortality for the tested doses.

## 1. Introduction

Entomopathogenic fungi (EPF) are important natural pathogens of arthropods. There are approximately 750 fungal species causing infection in insect and mite populations (Sinha et al., 2016). Fungi belonging to the order of Hypocreales, such as *Beauveria bassiana, Akanthomyces longisporum* and *Metarhizium anisoplae* have been well-studied and used widely as environmentally friendly biological control agents (Hesketh et al., 2008; Humber, 2008). Researchers in the field of biological control with EPF rely on robust results obtained via bioassays. A commonly used measure of efficacy when using infective propagules of microbial agents is lethal dose (LD) or lethal concentration (LC) (Finney, 1971). In order to obtain repeatable results when studying dose-time-mortality, it is imperative to conduct bioassays with a strict control of dosing method (Inglis et al., 2012). Infection of the insect host with fungal spores initiate via the cuticle (Ortiz-Urquiza and Keyhani, 2013), therefore the most common application of these microbial agents is topical. Depending on the size and number of insects and the target (habitat or insect surface), the method of application can vary greatly. The most frequently used method in applying fungal propagules on insect hosts is spraying (Inglis et al., 2012). There are a number of available devices including track- and handheld-sprayers designed for this purpose. The Potter spray tower is considered to be the standard of reference for such spraying techniques in the laboratory (Potter, 1952). However, this piece of equipment is stationary, takes up considerable laboratory space and is expensive. Hand-held sprayers are the cheapest option available, but they do not provide good control of the applied dose. Here, we provide a design for an appliance that could be readily made from inexpensive materials with relatively low cost. Similar setups are used frequently in laboratories working with microbial biocontrol agents, but the lack of published designs makes it difficult to construct robust, reproducible equipment kit for small scale spraying.

## 2. Methods

A portable micro-spray tower was designed following Mascarin et al. (2013) and Spence et al. (2020). The sprayer consists of an acrylic cylindrical tube, a top cap responsible for holding an artist airbrush (HP-SBS ECL3500 with standard nozzle, Iwata) securely in place with studded aluminium rods and a base cap acting as a sample tray. The airbrush is connected to a compressor (Powerjet Lite IS925, Iwata). The spray area allows space for a 90 mm Petri dish to be placed in the center (S1). The top cap with the fastening, the base cap and the feet (supplemental material) can be 3D printed or cut via a Computer Numeric Control (CNC) machine from a material of choice depending on the intended application. Here, we used 11.7 mm thickness PVC that can withstand repeated washing with 70% ethanol. The feet, base and top caps have 4 mm deep flanges connecting them to the acrylic tube. The parts are easily disassembled for cleaning and disinfection (S2).

Calibration of the device was carried out by spraying deionized water on 9 cm diameter filter paper (Qualitative 415, VWR). In order to understand the relationship between pressure, volume and droplet deposition, filter papers were weighted directly before and after spraying. Droplet deposition (µl/cm^2^) was calculated for all pressure and volume combinations of 8, 12, 16, 20, 24 PSI and 300, 400, 500, 750, 1000 µl resulting in 150 observations. A multiple linear model was used to describe the relationship between sprayed volume, pressure and liquid deposition on filterpaper.

The calibration was repeated with stain solution (Methyl blue, Sigma-Aldrich), for visual inspection of droplet deposition. Observations showed considerable fluctuations in pressure at 8 PSI and that aphids were flushed out of the petri dish at 24 PSI, thus these two pressure settings were excluded from further studies.

The relationship between fungal concentration and surface deposition of conidia was investigated using spore suspensions of *B*.*bassiana* 433.99^1^, *B*.*bassiana* 1787.18^1^ and *A. muscarius* 19.79^1^ at different concentrations ranging from 4.6×10^6^ to 3.7×10^8^ conidia/ml. Spore suspensions were sprayed on to 90 mm diameter Petri dishes containing 4-5 glass coverslips (22 × 22 mm). The applied volume was 400 µl at 12 PSI according to the previous study. All concentrations were tested at least three times. Sprayed cover slips were transferred to 45 ml centrifuge tubes containing 5 ml 0.01% Triton-X-100 and vortexed for 1 minute to dislodge conidia. Recovered conidial suspensions were enumerated using Fastread 102 (Biosigma) counting slides. Average number of conidia recorded from coverslips was used to quantify depositions rates (conidia/mm^2^). A linear regression model was used to describe the relationship between conidial deposition and concentration.

A dose response bioassay was carried out using *A. muscarius* Ve6 19-79 (Mycotal, NL) and even-aged apterous adult *Myzus persicae*. 18-20 aphids were treated in a 55 mm Petri dish with a concentration in the range of 1×10^5^-1×10^8^ conidia/ml. The seven tested concentration were replicated at least 3 times. Controls were treated with a sterile carrier (0.01% Triton-X-100). After treatment aphids were maintained on single leafs of Chinese cabbage (*Brassica pekinensis* var Wong Bok) in plastic cups with mesh covered vents at 20±1 °C and a 14:10 L:D regime. Aphid mortality was recorded daily for 7 days post-treatment. Nymphs were removed daily. Dead insects were surface sterilized with 70 % ethanol and rinsed in sterile distilled water. Sterilized cadavers were plated separately and observed for fungal outgrowth to confirm that death was associated with fungal infection.

## 3. Results

Droplet deposition showed the least spread (2.62 ± 0.01 µl) with a volume of 400 µl sprayed at 12 PSI. A significant relationship between water deposition, volume and pressure was indicated by the multiple linear model (F_2, 147_=5522; P<0.0001; adjusted R^2^=0.989) (Figure 1).

**Figure 1.**
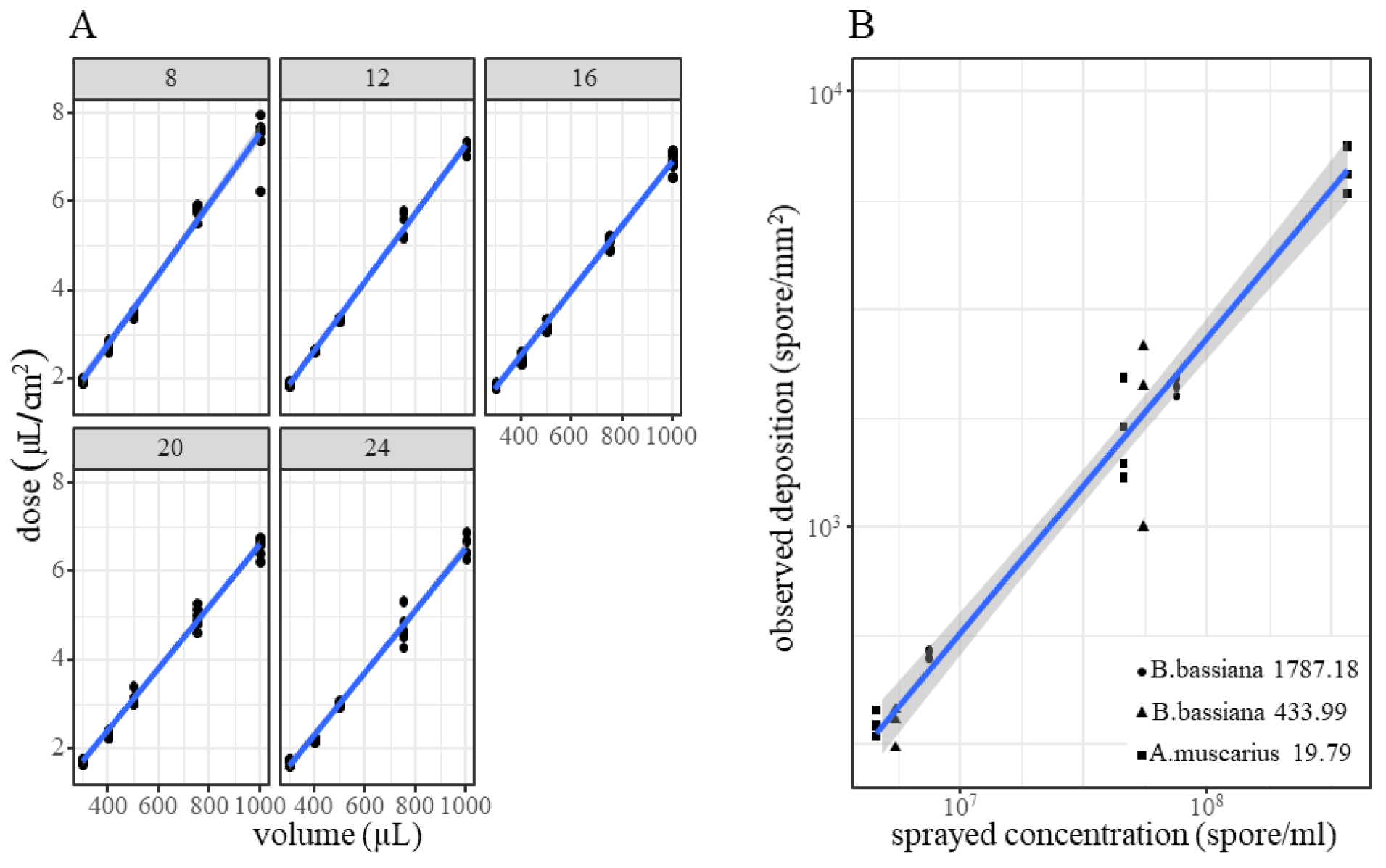
Water deposition expressed as dose at different applied pressures and sample volumes. Panel headings (8-24) refer to application pressure in PSI (A). Observed conidia deposition at different applied concentrations (B).

Visual inspection of spray patterns on the filter papers indicated that 400 µl sprayed at a pressure of 12 PSI provided the most even coverage.

The determination of spore deposition was limited to the range where spore counts could be accurately enumerated by the counting slides. There is a significant relationship between conidia deposition and applied concentration (F_1, 22_=449.3; P<0.0001; adjusted R^2^=0.951). The results show that within concentration variation is low, which is key for dose-response testing of microbial propagules (Figure 1).

The dose-response bioassay resulted in repeatable mortality for doses spread in a wide range (Figure 2). The variation within doses could be attributed to the differences in aphid batches and the physiology of cabbage leafs cut from plants.

**Figure 2.**
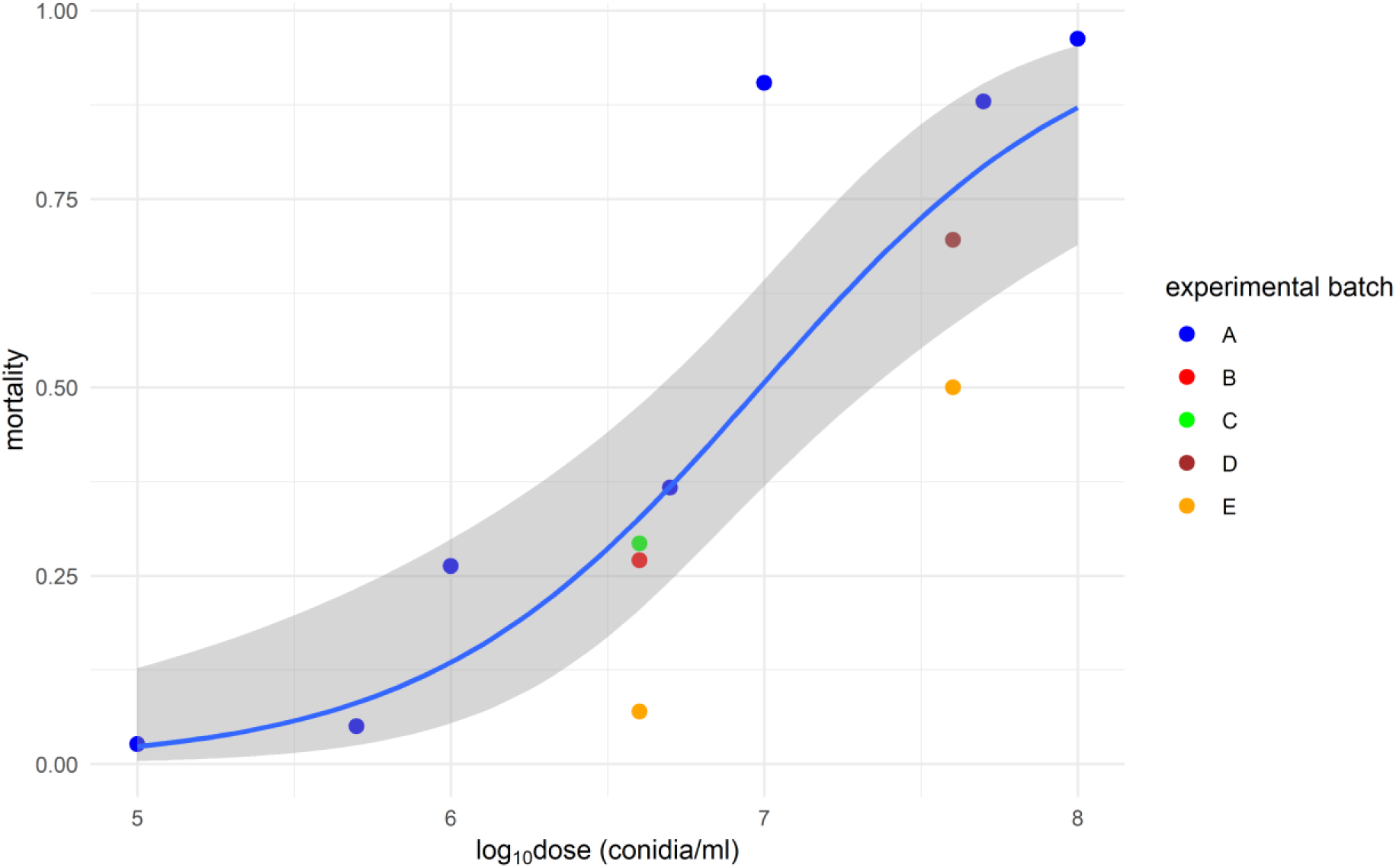
*M. persicae* proportion mortality after 6 days of exposure to *A. muscarius* depending on tested concentrations. Points represent pooled cumulative mortality recorded from individual aphid cups. Colours represent different aphid batches (blocks). A quasibinomial regression model (blue line) was fitted to predict the dose response. Grey area represents 95% confidence intervals.

## 4. Discussion

The described device provides a relatively cheap alternative to the commonly used laboratory spraying equipment in foliar testing of microbial agents or pesticides. In comparison to the Potter spray tower, the advantages of the design includes inexpensive spare parts, portability, and usage in laboratories with restricted space. The most expensive parts of the presented setup are the compressor and the airbrush. The cost of the acrylic tube and the PVC sheet including the CNC machining of tower parts (four replicates) was under $600. These costs could be further reduced by material selection. The specifications of the tower (height, diameter) could be easily adjusted to suit individual preferences. The airbrush used here has numerous feed and nozzle attachements that can hold higher load volumes and produce different spray patterns (https://www.iwata-airbrush.com). However, the tower parts can fit, or can be modified to fit other similar airbrushes. Limitations related to the range of applied pressure was observed due to the low weight of aphids tested in our bioassays. Such constraints should not arise with heavier targets or applications conducted on leaf.

The design described here can be adjusted to accomodate the needs of most laboratories aiming to conduct microbial agent or pesticide testing without investing in expensive kit or restricting considerable space for stationary equipment. Further improvements were made by attaching a vacuum filter, to reduce contamination when opening the chamber. Alternatively the sprayer can be operated in a fumehood. The Computer Aided Design (CAD) files that can be used for 3D printing, CNC routing or modification to user requirements are available as supplementary materials.

## Supporting information

design files

## Acknowledgments

We are grateful to Professor David Chandler, Dr. Gillian Prince for the fungal strains and providing useful discussion and comments. We thank Dr. Mark Mallott for the photographs. This work was supported by the Agricultural and Horticultural Development Board (CP 176).

## Footnotes

1 Warwick Crop Centre Culture Collection strain number

## Notes

### Competing Interest Statement

The authors have declared no competing interest.

### Summary of Updates

Supplemental files updated.

