## Supplementary figures and images for "Laboratory sprayer for testing of microbial biocontrol agents: design and calibration"

### S1

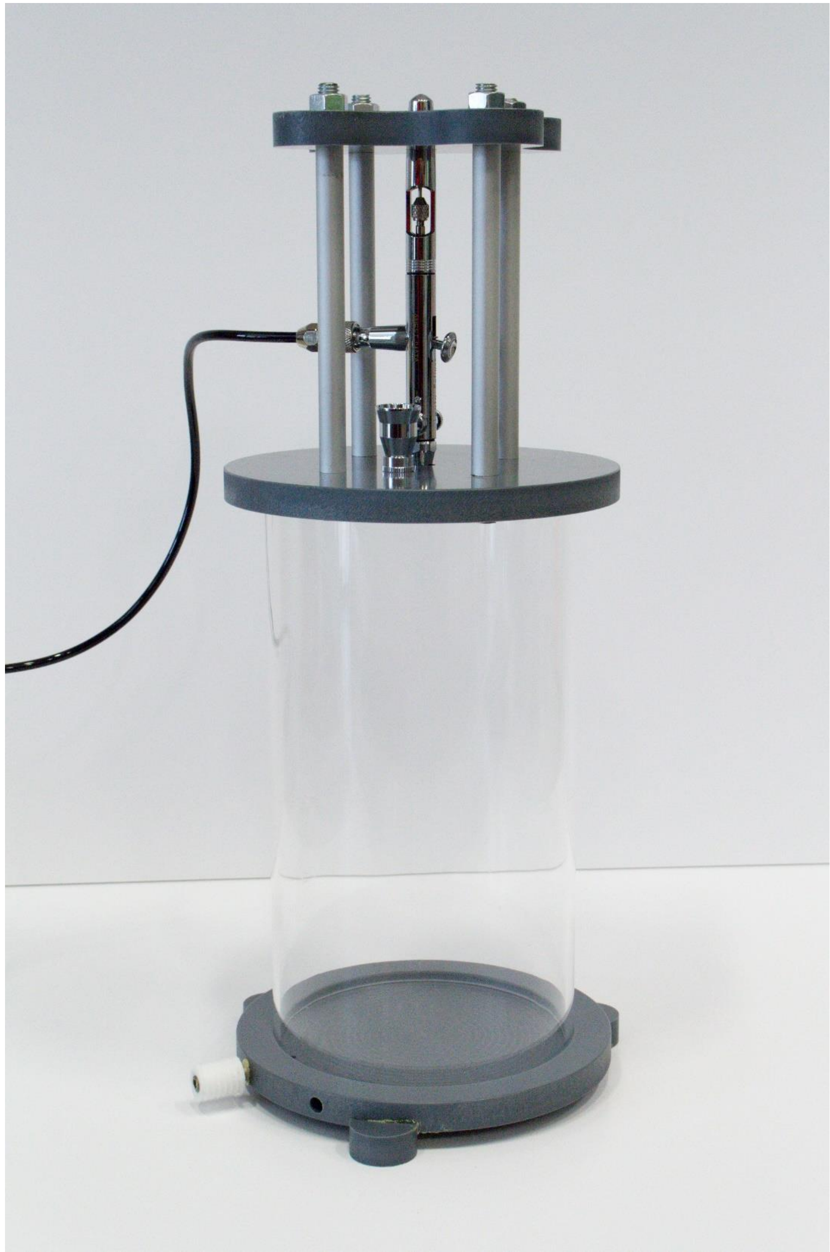

S1 Spray tower assembly

### S2

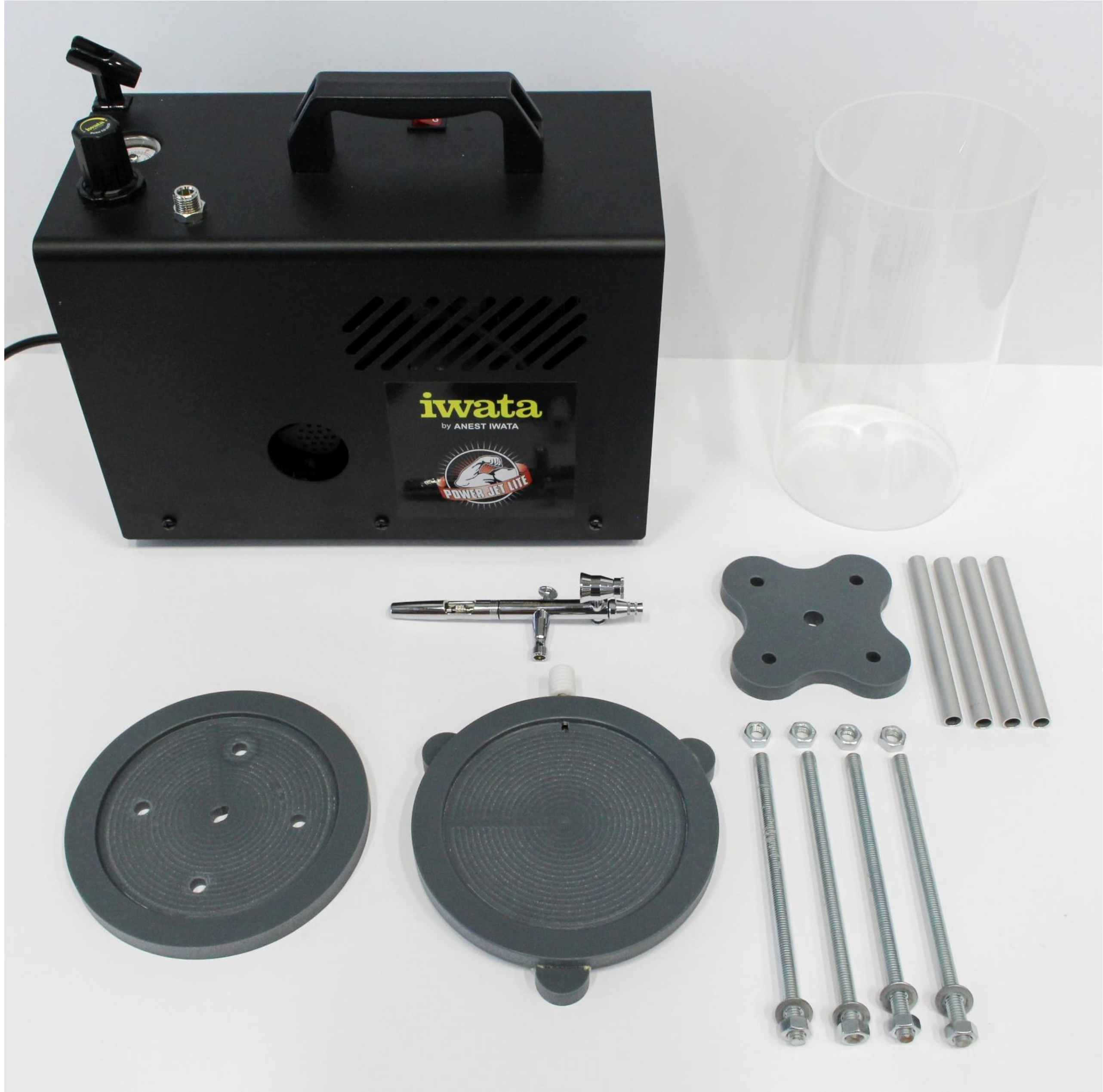


S2 Spray tower parts
