## Supplementary material for "Laboratory sprayer for testing of microbial biocontrol agents: design and calibration": design files: Spray tower annotated2.pptx

### Slide 1
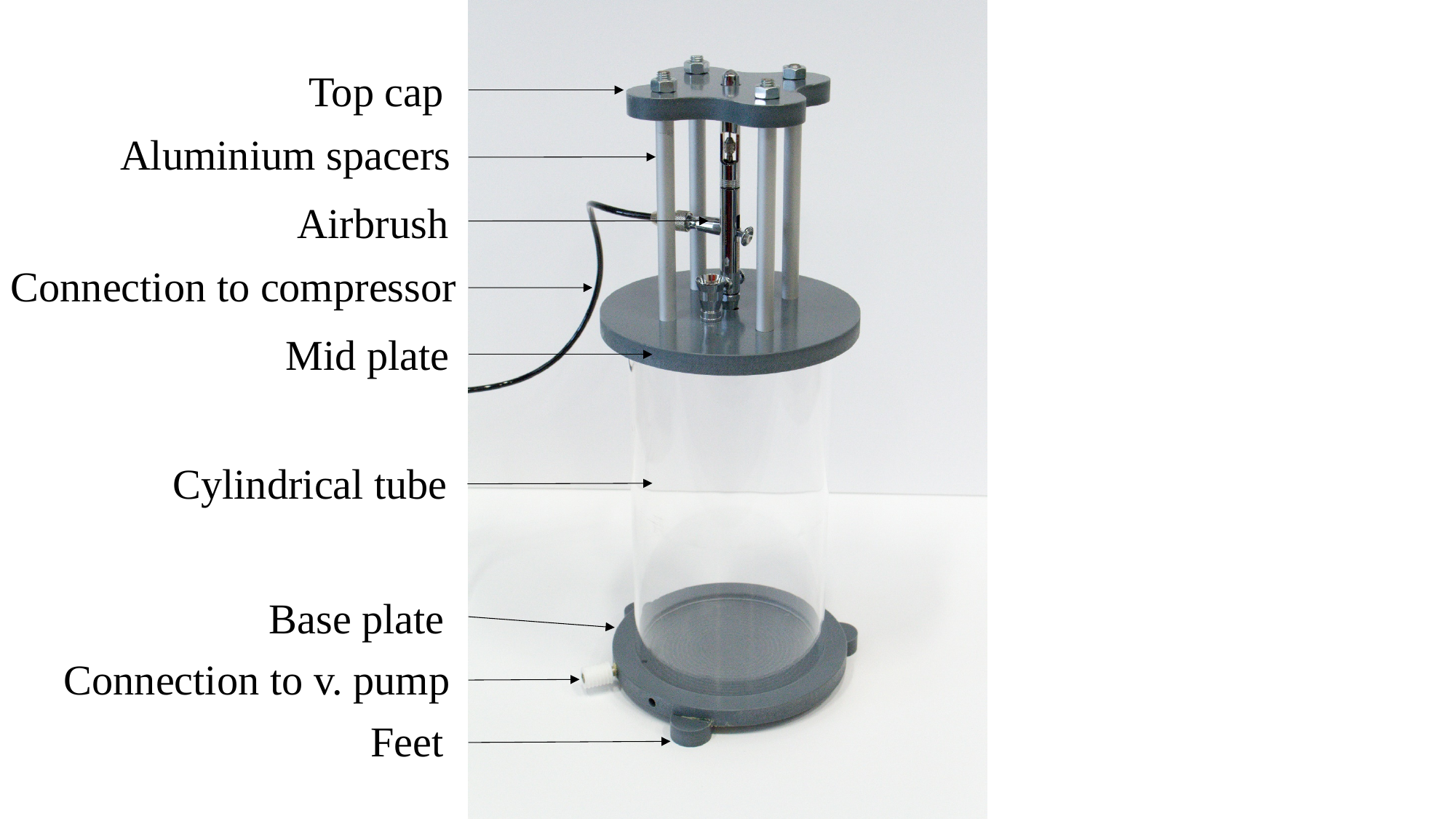

Top cap
Aluminium spacers
Airbrush
Connection to compressor
Mid plate
Cylindrical tube
Base plate
Connection to v. pump
Feet
